# Audible Feedback Improves Internal Model Strength and Performance of Myoelectric Prosthesis Control

**DOI:** 10.1101/259754

**Authors:** Ahmed W. Shehata, Erik J. Scheme, Jonathon W. Sensinger

**Affiliations:** University of New Brunswick, Electrical and Computer Engineering, Fredericton, E3B5A3, Canada; Institute of Biomedical Engineering, Fredericton, E3B5A3, Canada

## Abstract

Myoelectric prosthetic devices are commonly used to help upper limb amputees perform activities of daily living, however amputees still lack the sensory feedback required to facilitate reliable and precise control. Augmented feedback may play an important role in affecting both short-term performance, through real-time regulation, and long-term performance, through the development of stronger internal models. In this work, we investigate the potential tradeoff between controllers that enable better short-term performance and those that provide sufficient feedback to develop a strong internal model. We hypothesize that augmented feedback may be used to mitigate this tradeoff, ultimately improving both short and long-term control. We used psychometric measures to assess the internal model developed while using a filtered myoelectric controller with augmented audio feedback, imitating classification-based control but with augmented regression-based feedback. In addition, we evaluated the short-term performance using a multi degree-of-freedom constrained-time target acquisition task. Results obtained from 24 able-bodied subjects show that an augmented feedback control strategy using audio cues enables the development of a stronger internal model than the filtered control with filtered feedback, and significantly better path efficiency than both raw and filtered control strategies. These results suggest that the use of augmented feedback control strategies may improve both short-term and long-term performance.

## Introduction

Recent advances in material design, micromachining, and the understanding of human neuromuscular systems have enabled the development of lightweight prosthetic devices that can be used to help amputees perform activities of daily living. One approach to controlling these devices is to use myoelectric signals sensed from contractions of the amputee’s remnant or congenitally different muscles^1^. Researchers have developed many signal processing techniques^2^, feature extraction methods^3–5^, and control strategies^6^ to enhance the performance of this approach. Despite these advancements in the field of myoelectric prostheses, many amputees still abandon their devices out of frustration^7^, due in part to insufficient precision in the control of prosthesis movements and a lack of adequate sensory feedback^8,9^.

Although invasive feedback, such as stimulation of sensory peripheral nerves^10^, has the potential to elicit close-to-natural tactile sensations, many prosthesis users prefer non-invasive feedback methods that do not require surgical intervention^11^. With this preference in mind, researchers have proposed using non-invasive sensory substitution methods to provide sensory information to prostheses users either through different sensory channels or using different modalities^12^. Vibro-tactile^13^, mechano-tactile^14^, electrotactile^15–17^, skin stretch^18^, and auditory^19^ are just some of the techniques that have been developed and used to provide prosthesis users with feedback. Although some studies (e.g.,^14,20,21^) have shown that sensory feedback improves performance, others *(c.f*.,^14^) have concluded that sensory feedback had no effect on performance. This lack of consensus arises, at least in part, because of an unclear understanding of how the incorporation of feedback relates to performance.

The role of feedback for real-time regulation and in improving human understanding of the control system and the task being performed is still unclear. Researchers have hypothesized that the human central nervous system implements control by estimating the current state of the musculoskeletal system and updating this estimate using sensory feedback^22,23^. This state estimation and prediction process is governed by a model formed in the central nervous system, which is known as internal model. This internal model holds properties of the arm, which are used to imitate its behavior^24^, predicts consequences of an action, and computes an action based on desired consequences^25^. Hence, this internal model is used in the feedforward control^26^ of the arm or residual limb, and affects the overall performance of a prosthesis^27^.

Humans use visual feedback along with other sensory and proprioceptive information to develop their internal model, however constant visual attention and the high level of concentration it requires may lead powered prosthesis users to reject their devices^28^. Conveniently, augmented feedback can be used to convey artificial proprioceptive and exteroceptive information^21^’^29^, which may help to develop strong internal models. Researchers have used audio augmented feedback in both robotic teleoperation^30,31^ and Brain Computer Interfaces (BCI)^32^ and have concluded that audio augmented feedback improves performance. Unlike visual feedback, audio requires less focus of attention and reduces distraction^33,34^. We hypothesize that audio augmented feedback may similarly improve the performance of myoelectric prosthesis control by enabling the development of stronger internal models.

In a recent study^35^, we assessed the performance and the strength of internal models developed by users when using two myoelectric control strategies that differed in control signals as well as level of feedback. The main aim of that study was to investigate whether the ability of users to adapt is influenced by the degree of sensory feedback inherent in different myoelectric controllers. Results from that study suggest that control strategies with raw control signals and high feedback level (Raw Control with Raw Feedback (RAW), such as with regression-based control) enable the development of a strong internal model, but at the expense of short-term performance. In addition, we found that control strategies with filtered control signals and reduced feedback (Filtered Control with Filtered Feedback (FLT), such as classification-based control) may enable better short-term performance, but hinder the development of internal models.

To mitigate this tradeoff, we have extended this work by decoupling the concepts of control and feedback through the use of augmented feedback. We combined the filtered control strategy that resulted in better short-term performance with audio augmented feedback from the raw control strategy that enables the development of stronger internal models. In this work, we assessed internal model strength and short-term performance for this audio augmented feedback control strategy known as Filtered Control with audio Augmented Feedback (AUG) and compared its results to the two commonly used myoelectric control strategies, RAW and FLT, that we assessed previously^35^. Our results show that the audio augmented feedback control strategy produces better short-term performance (as assessed using path efficiency and accuracy) than the feedback-rich control strategy, while enabling the development of a stronger internal model than the reduced feedback control strategy.

## Results

To inform the development of myoelectric control strategies with better short- and long-term performance, we investigated whether augmented feedback could improve user’s internal model strength without reducing short-term performance of the control. Short-term performance and internal model strength (as a predictor for long-term performance^36^) were evaluated while using a filtered control strategy that was augmented by audio feedback in a virtual target acquisition constrained-time task.

### Outcome Measures

To assess the internal model developed by the human central nervous system and evaluate performance of the audio-augmented controller, we employed the same five parameters that were used in our previous work^35^:

- Adaptation rate was extracted by computing the rate of feedforward modification of the control signal from one trial to the next^37,38^. The first 100 – 230 msec window of activations that started on the mark of the visual rendering on the screen for each trial was used to ensure that only the subject’s feedforward intent was captured and to avoid the effect of feedback synchronization. The target control signal was the activation of the cursor movement to the right direction only i.e., wrist extension. Other activations were considered as self-generated errors, which subjects were instructed to minimize.
- The just-noticeable-difference (JND) parameter was computed from a perception threshold test as the threshold value reached after 23 reversals calculated by an adaptive staircase^39^. This parameter is influenced by both controller and sensory noise.
- Internal model uncertainty *P_param_* was calculated using sensory uncertainty, controller uncertainty, JND, and adaptation rate^35^.
- Performance of the audio-augmented control strategy was evaluated using path efficiency, computed by comparing the path taken to reach a target to the shortest Manhattan path to that target. Targets were defined by their location with respect to the X and Y axes on the computer screen. Targets that were located on either the X-axis or the Y-axis were known as on-axis targets, which only required the activation of a single DOF to be acquired. By the same token, targets that were not located on either the X-axis or the Y-axis were known as off-axis targets and therefore requiring the activation of 2 DOFs to be acquired (Appendix Figure A1).
- Accuracy of target acquisition was computed as the Manhattan distance between the center point of a target and the actual final point reached. The ratio between this value and the optimal Manhattan path was then used to compute normalized accuracy as a percentage.

### Psychophysical tests

The adaptation rate serves as an indicator of how much humans modify their internal models when performing a task. A value of 1 indicates perfect adaptation; higher and lower values correspond to over- and under-compensation, respectively. Results for the adaptation rate test show that there was a statistically significant difference in adaptation rates to self-generated errors between subjects when using the RAW, FLT, and AUG controllers, as determined by a one-way ANOVA (F (2, 21) = 9.17, p = 0.02). A Tukey HSD post hoc test reveals that the adaptation rate for subjects using FLT (0.46 ± 0.19) was significantly lower than RAW (0.98 ± 0.3, p = 0.02) and AUG (0.86 ± 0.27, p = 0.021). In addition, no significant difference was found between adaptation rate data for subjects using the RAW and AUG control schemes (p = 0.65). These results suggest that audio-augmented feedback may enable similar adaptation behavior to feedback-rich control strategies (Figure 1. a).

**Figure 1.**
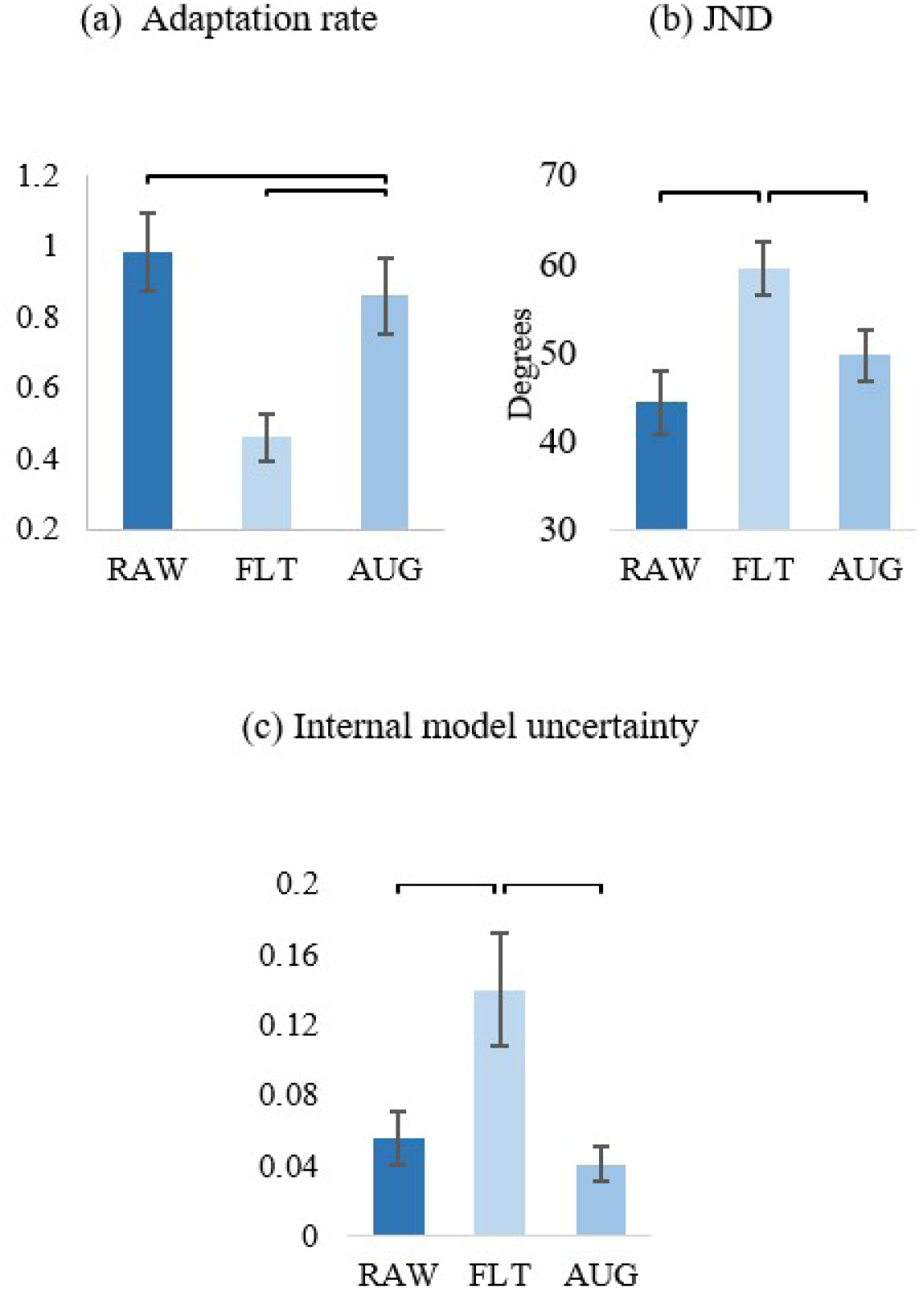
Overall psychophysical test results. (a) Results for adaptation rate across control strategies showing low adaptation rate to self-generated errors when using FLT. (b) JND results across control strategies showing low perceivable sensory threshold when using either RAW or AUG. (c) Internal model uncertainty reflected by both adaptation rate and JND results showing significantly less uncertain (more confident) internal models developed when using RAW and AUG than the one developed when using FLT.

The JND parameter is a measure of the smallest measure of stimulus that a subject is able to identify when using a certain controller. The lower this parameter, the better the ability of a subject to detect changes in the controller used. Like the adaptation rate results, analysis of JND test data using one-way ANOVA revealed a significant difference between subjects using RAW, FLT, and AUG (F (2, 21) = 5.17, p = 0.008). Upon running a Tukey HSD post hoc test on the data, we found that subjects who tested FLT had significantly higher JND values (59 ± 16 degrees) than other subjects who tested RAW (45 ± 20 degrees) and AUG (47 ± 13 degrees), but no significant difference in JND data between subjects who tested RAW or AUG (p = 0.92), which suggests that feedback in both RAW and AUG was sufficient to allow subjects to perceive a lower sensory threshold than the reduced feedback FLT (Figure 1. b).

Both adaptation rate and JND results were used to compute the internal model uncertainty for each of the tested control strategies. The lower the internal model uncertainty, the more confident a subject is in the control system used. Results showed significant difference in internal model uncertainty between the control strategies (robust Welch ANOVA (F (2, 13) = 9.3, p = 0.003)). Games-Howell post hoc analysis revealed a significant difference between internal model uncertainty for subjects who tested FLT (0.14 ± 0.1) and the subjects that tested RAW or AUG (0.055 ± 0.09, p = 0.035 and 0.041 ± 0.034, p = 0.004, respectively). On the other hand, there was no significant difference in the internal model uncertainty between subjects that used RAW or AUG (p = 0.56) (see Figure 1. c). These findings suggest that a strong internal model may be developed when using either the RAW or AUG control strategy, which confirms the first part of our hypothesis that audio-augmented feedback may enable the development of a stronger internal model than FLT alone.

### Performance tests

For on-axis path efficiency, there was a significant difference between control strategies as determined by a one-way ANOVA (F (2, 21) = 5.8, p = 0.01). Bonferroni post hoc analysis showed that subjects who tested the AUG control strategy had significantly higher path efficiency than subjects who tested either the FLT or RAW strategies (p = 0.019, p = 0.02) (Figure 2. a). Analysis of the accuracy data collected from subjects using the three control strategies showed a significant difference between control strategies (one-way ANOVA (F (2, 21) = 4.5, p = 0.024)). Even though subjects that used FLT achieved high accuracy for on-axis targets (73 ± 7%), it was not significantly different (Bonferroni post hoc test) from subjects using who used RAW (65 ± 11%, p = 0.36) or AUG (80 ± 11%, p = 0.51). Conversely, subjects who tested AUG achieved significantly higher accuracies than those who tested RAW (p = 0.02) (Figure 2. b). These results further support our hypothesis that augmented feedback may also enable better short-term performance. For off-axis target performance tests, the null-hypothesis was not rejected when comparing path efficiency or accuracy data between the three control strategies (one-way ANOVA (F (2, 21) = 0.16, p = 0.86)) (Figure 2. c and d).

**Figure 2.**
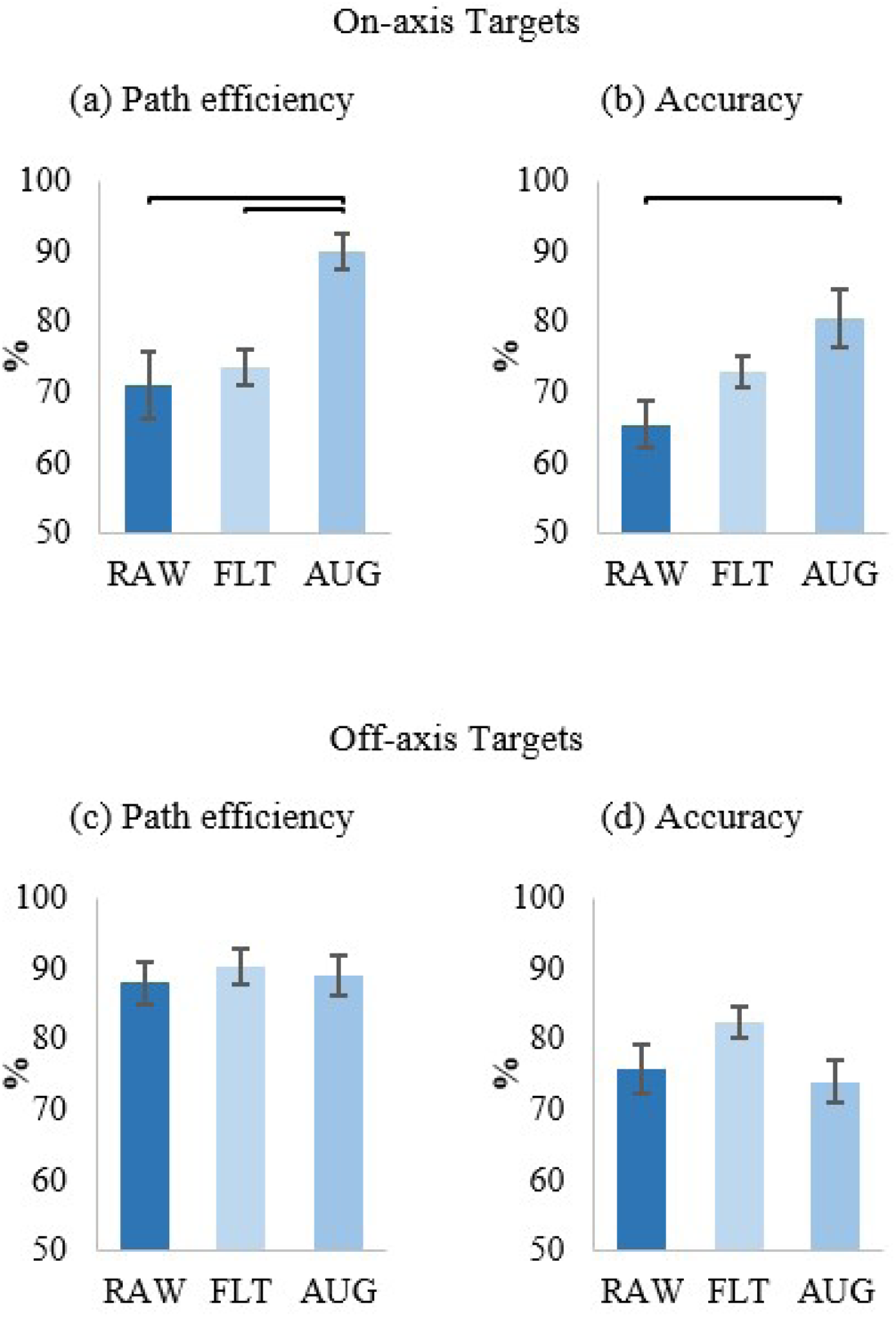
Overall performance results. (a) and (b) show results for path efficiency and accuracy, respectively, of on-axis targets across control strategies. (c) and (d) show no significant difference between control strategies tested for off-axis targets.

### Learning effect

We ran 28 dependent paired t-tests to investigate possible learning effects when testing with one control strategy and retesting using another one (summarized in Appendix). Unlike adaptation rate, a significant learning effect was observed for JND values of subjects who used FLT after being exposed to AUG, where the JND dropped from 74 ± 8.0 degrees to 43 ± 4.2 degrees (p = 0.029). This suggests that there may be possible enhancement in sensory perception threshold when using the FLT control strategy after first being exposed to the feedback-richer AUG control strategy. On the other hand, learning effect for JND values for subjects who tested FLT after AUG were not significantly reflected in the internal model uncertainty (t (3) = 1.41, p = 0.25), because internal model uncertainty is affected by several other parameters e.g., adaptation rate and controller noise. These results suggest that internal model uncertainty for FLT may be improved after being first exposed to AUG by improving sensory perception, and less so by adaptation. Interestingly, on-axis target path efficiency significantly increased when testing the AUG control strategy after testing the feedback-rich RAW control strategy (t (3) = -14, p = 0.005). Also, accuracy for on-axis targets when using FLT after first being exposed to AUG increased significantly from 84 ± 3.7% to 94 ± 3.1% (t (3) = 23, p = 0.028).

## Discussion

Significant progress made in the fields of signal processing, sensory substitution and pattern recognition for myoelectric prostheses has allowed for improvements in the performance of myoelectric control. It stands to reason that such improvement is partially driven by feedback^40^, however contradictory results about the effect of feedback (in presence of vision) on performance oppose this belief^41^.

Several studies have investigated the effect of various feedback modalities on performance^12^. In particular, results from studies providing electromyography (EMG) biofeedback either through visual^42^ or electrotactile^20^ feedback have showed promising improvement in performance, but have not investigated the effect of this feedback on the internal model. Other researchers^36^ have investigated the effect of internal models on performance and found evidence of improvement in performance through improvement in the internal model. In a recent study^43^, researchers have found that feedback improves short and long-term performance and credited this improvement to internal models. However, they were not able to quantify the improvement in internal models that influenced the improvement in performance. To address this deficiency, in a previous study^35^ we used a psychophysical framework to quantify the strength of internal models developed for two commonly used myoelectric prosthesis controllers^44^, namely regression-based and classification-based controllers. In this study, we extended these previous studies by exploring the effect of relaying EMG biofeedback through the less-focus demanding audio feedback to augment myoelectric controllers that enable better short term performance. We hypothesized that augmented feedback may enable stronger internal model generation and better short-term performance. Our results showed that an audio-augmented feedback control strategy enables the development of a significantly stronger internal model than a classification-like filtered control with filtered feedback control strategy. This audio-augmented feedback controller also resulted in significantly better path efficiency for a one DOF task than both the raw feedback and the filtered feedback controllers.

Unlike classification-based control methods, the use of a regression-based control strategy enables the extraction of more information that may be used for augmented feedback. In this work, the concept of a hybrid control strategy that combines the robustness of classification-based control and the rich feedback of regression-based control was introduced. The concepts of feedback and control were effectively decoupled by employing classifier-like control, while providing the user with audio-augmented feedback derived from the feedback-rich regression control scheme.

Adaptation rate results showed that the controller with audio-augmented feedback enabled better adaptation behavior than FLT while maintaining similar adaptation behavior to the feedback-rich RAW. Hence, we may conclude that audio-augmented feedback may contain sufficient and comparable information as the RAW, therefore enabling better understanding of the myoelectric system than FLT. Equally as important, JND results for sensory perception showed that subjects who used either RAW or AUG achieved lower sensory thresholds than those who used FLT, which again supports the conclusion that feedback inherent in AUG was as effective as the feedback in the feedback rich RAW control strategy, enabling the perception of lower sensory thresholds. These results combined with results from the assessment of internal model uncertainty confirm our hypothesis that audio-augmented feedback enables the development of a strong internal model, possibly leading to better long-term myoelectric control performance.

On-axis performance results showed that audio-augmented feedback helped subjects perform better with respect to path efficiency and accuracy when compared to FLT. This finding suggests that subjects are able to integrate the audio feedback and use it for real time regulation as well as for internal model development. We also found that when subjects were first trained with AUG and then used FLT, they retained the internal model developed for AUG and were able to achieve better on-axis accuracy. This observation raises a new question to be investigated in future work: how long can this stronger internal model be retained before reverting completely to a model developed by the FLT control strategy?

Despite significant improvement in performance for on-axis targets when using the audio augmented controller, no significant improvement for off-axis targets accuracy or path efficiency was found. However, statistically significant results for accuracy and path efficiency for off-axis targets may be found with larger cohorts targeted towards these metrics. Future work informed by this study could include investigating the effect of augmented feedback on the internal model strength and short-term performance when using other controllers, assessing the impact of other modalities of augmented feedback (such as vibrotactile), exploring combinations of augmented feedback (such as audio + vibrotactile, which might enable an even stronger internal model), investigating how long the human central nervous system is able to retain the improved internal model strength (following a protocol similar to other studies^43^), and finally, assessing the benefits of using augmented feedback for limb-different individuals by performing psychophysical tests using prosthetic devices. Future studies can extend this work to accommodate the integration of feedback in real-time regulation using a framework such as the Smith predictor^45,46^ or the adaptive Smith predictor^26,47^.

Some limitations to this study were: the learning effect due to prolonged exposure to the feedback-rich control strategy was not investigated, no amputees were included in this study, and only a one DOF task was used for the assessment of the internal models.

The benefits of using audio-augmented feedback for myoelectric prosthesis control, highlighted in this study, call for inclusion of audio feedback in myoelectric control training procedures. For example, myoelectric prosthesis users may be provided with audio feedback when training with a myoelectric controller to improve their internal model strength and then prompted to use this control strategy without this audio feedback during regular use. One might argue that providing audio feedback using an earpiece or headphones to prostheses users might be impractical or uncomfortable, but emerging technologies, such as those that transmit sound using bone conduction, may soon enable users to benefit from other types of similarly augmented feedback. Furthermore, it is not yet known how frequently this augmented feedback is need. It is possible that users could occasionally enable the audio-augmented feedback to re-enforce their understanding.

## Methods

### Control Strategies

In order to conduct a fair comparison between the control strategy developed here and the ones assessed in^35^, Support Vector Regression (SVR) was selected as the machine learning algorithm for the AUG strategy^48^. Figure 3 shows a sample of raw signals sensed from muscles. These signals were processed using standard EMG processing techniques used in the field of prostheses^49,50^. Specifically, these signals were low-pass filtered at 450 Hz with a fifth-order Butterworth filter and high-pass filtered using a third-order filter at 20 Hz^51^. In addition, a notch filter was implemented using a 2nd-order Butterworth band-stop filter from 57 Hz to 63 Hz^52^. Data were sampled at a frequency of 1 KHz and time-domain features^53,54^ were extracted from these processed signals every 64 msec using 160 msec windows to train the SVR model.

**Figure 3.**
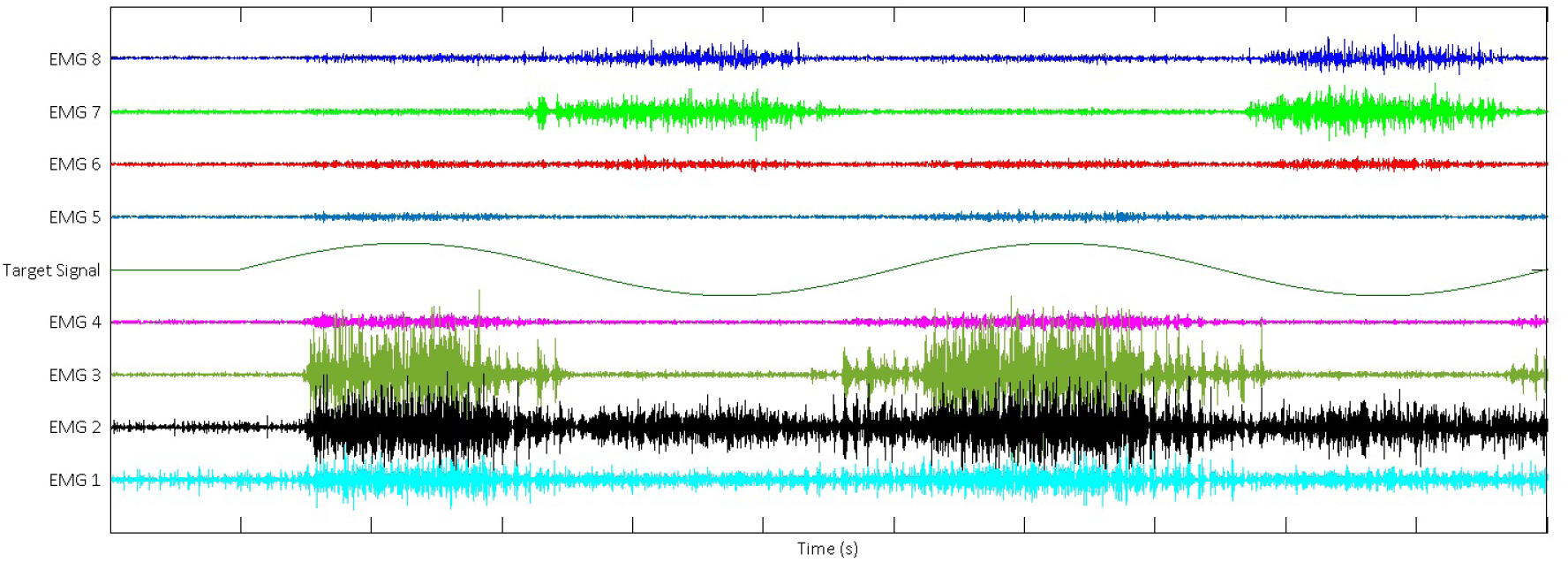
A sample of EMG signal recorded from 8 electrodes placed on subject’s forearm. Subjects were asked to follow a cursor on a screen moving to the left and right with isometric wrist extension/flexion. The velocity profile for the moving cursor is shown in the middle of this figure as a sinusoidal wave with a 1 second delay after which the cursor started moving to the right first.

For real-time control, the SVR model produced output every 16 msec using 160 msec windows^51^. The output from this model was directly mapped to the velocity of the cursor for simultaneous activation of 2 degrees-of-freedom (wrist extension/flexion and adduction/abduction), but only the highest activated DOF was used in FLT; therefore, only allowing sequential control. Similar to FLT, the AUG control strategy only allowed sequential control, but used audio feedback to convey simultaneous control information, making this controller a hybrid between classification-based control and regression feedback (Figure 4).

**Figure 4.**
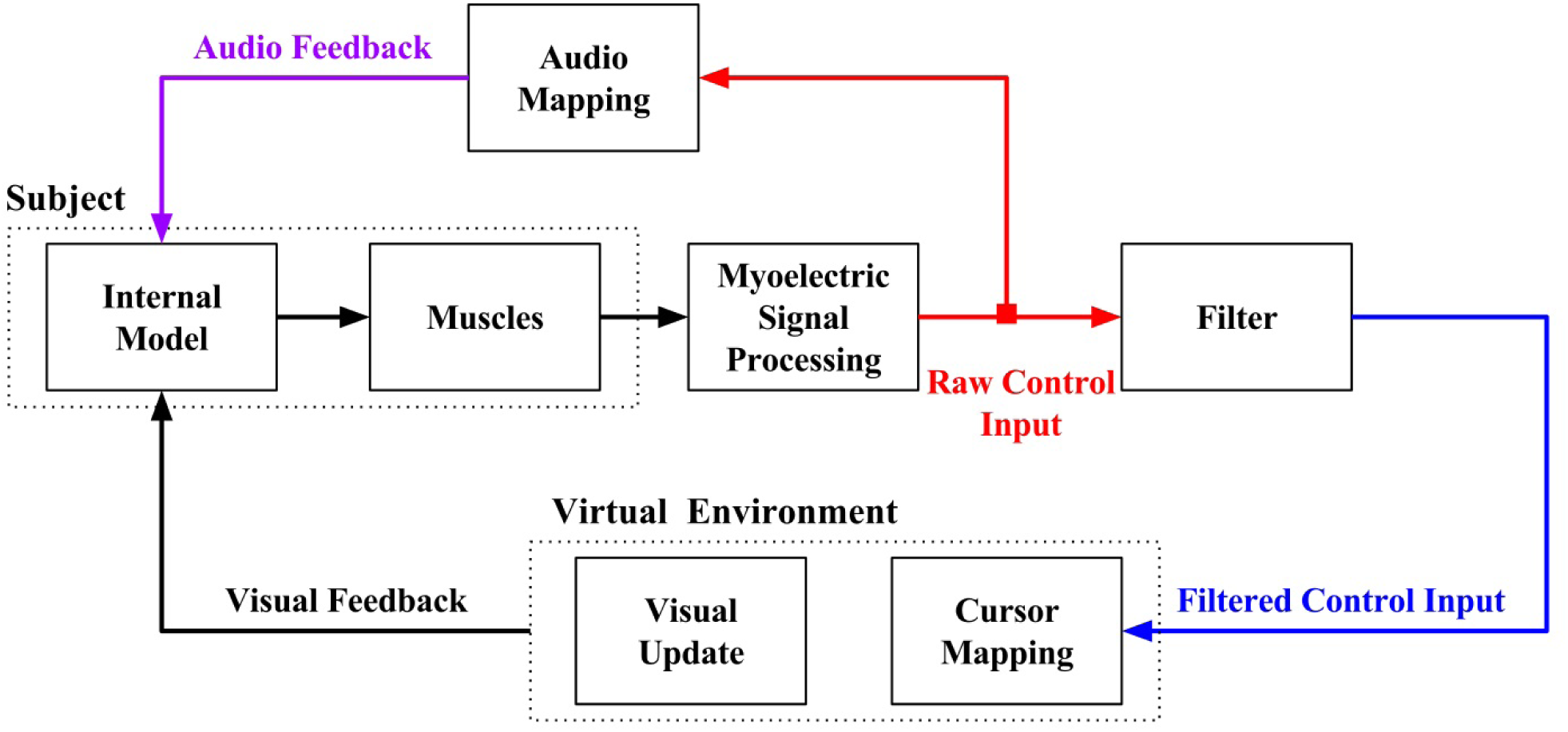
Control strategy development. Audio feedback mapped from raw control input and augmenting the filtered control input.

### Audio Augmented Feedback

For the AUG control strategy, audio feedback maps the simultaneous activation of the 2 DOF motion using 4 audibly distinct frequencies i.e., one frequency per direction, (see Figure 5.a). The amplitude of the tone is directly proportional to the SVR output, requiring no knowledge of the intended task. For instance, a subject using AUG to reach a point to the right of the cursor while activating both the right and up directions, but with a higher activation in the right direction will see the cursor moving to the right accompanying to that cursor movement, the subject will hear two tones simultaneously with frequencies of 500 Hz and 800 Hz with the 500 Hz tone being louder than the 800 Hz tone.

**Figure 5.**
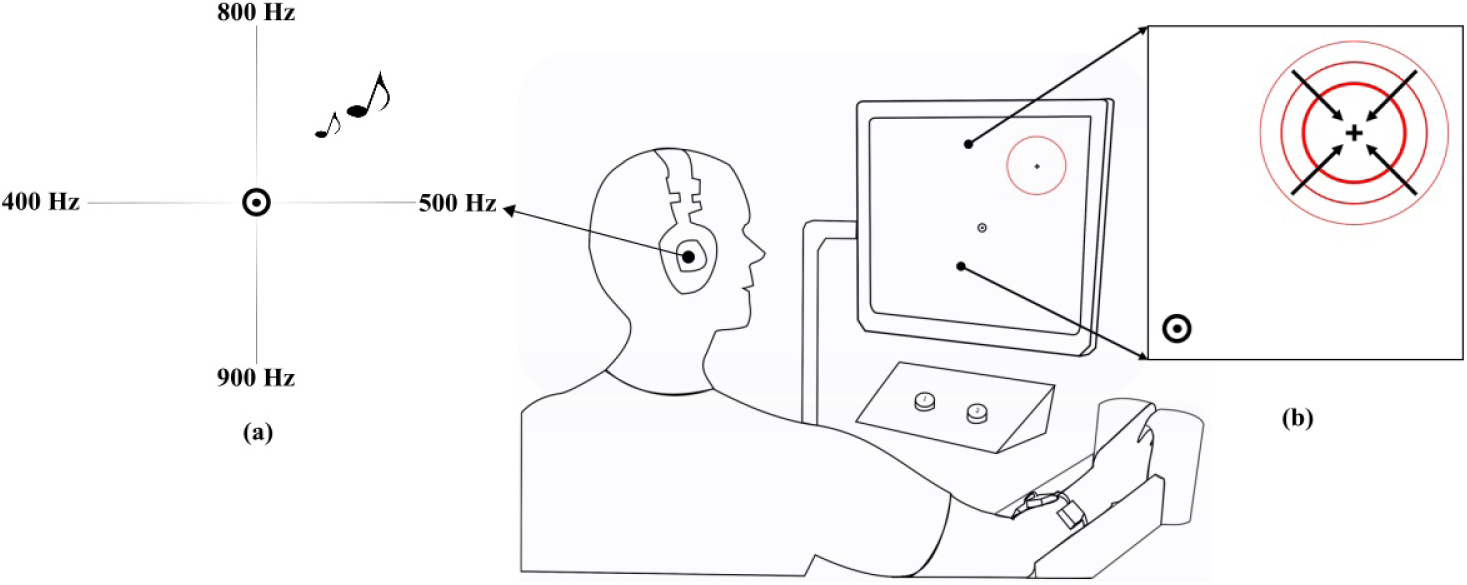
Experimental Set-up, with a subject controlling a cursor on the screen to acquire a set of targets during the training phase. The target shown on the f Subjects were allowed to make their choice during the just-noticeable-difference test by pressing keys on a keyboard.(a) Audio map showing the frequencies assigned for each direction, i.e., 500/400 Hz corresponded to wrist extension/flexion and 900/800 Hz corresponded to wrist adduction/abduction. The volume is directly proportional to the magnitude of the activation in each direction (b) A shrinking off-axis target.

### General Experimental Procedures and Protocol

Each subject sat comfortably in a chair approximately 80 cm away from a computer screen with their right arm relaxed in a restraint and used their left hand to press keys on a keyboard placed on a desk in front of them. Subjects were asked to wear a 15 mW Cobra headset with the volume set to a no more than 52.5 ± 3 dB for the whole period of the experiment, but were allowed to remove it during scheduled breaks between the testing block. During the testing blocks, subjects controlled a cursor on a computer screen using isometric muscle contractions sensed by an electrode array placed on their forearm to acquire targets (Figure 5.b). Targets were crosshairs surrounded by an imploding red circle. Using the same experimental protocol implemented in^35^, which was approved by the University of New Brunswick’s Research and Ethics Board (file number 2014–019), subjects used each control strategy to complete a series of test blocks in a specific order after accomplishing a training block.

To determine the gain for mapping the control signals to the cursor velocity on the computer screen, each subject was asked to control a brush to paint a screen^35^. Their ability to maximize the area covered by paint determined the appropriate gains for each DOF, i.e., 20 pixels/s for wrist adduction/abduction DOF. The training block consisted of three sets of targets that appeared at predetermined, but randomly ordered, positions. Each set consisted of 16 targets, which subjects were asked to acquire in less than 12 seconds, after which it would disappear and another target would appear. The first test block was used to test the adaptation to self-generated errors. In this block, subjects were asked to repeatedly acquire a single target on the horizontal axis over a set of 80 trials.

Before the start of each test block, subjects were given a 2 minutes break, in which they were allowed to stand, remove the headset, and stretch if needed. The JND perception threshold test, the second test block, is a psychometric measure of sensory threshold for perception of a sensory stimulus. In this test, subjects were presented with two trials where one of these trials was perturbed with a stimulus. Subjects were then asked to select the trial which they thought had the added stimulus by pressing 1 for trial 1 and 2 for trial 2 on a keyboard^39^. The added stimulus was calculated using an adaptive staircase with accuracy set to 0.84^55^. Finally, the last test block was a performance test where subjects were asked to acquire targets, arranged in 2 sets of 16 targets each. For the first set of targets, subjects were tasked with acquiring each target in less than 1.7 s, whereas they were only allowed 1.4 s for each target in the second target set.

### Participants

A total of twenty-four healthy right hand dominant subjects (15 male, and 9 female; mean and SD of age: 25 ± 6 years) participated in the study. All participants had either normal or corrected-to-normal vision. Informed consent was obtained from subjects before conducting the experiment according to the University of New Brunswick’s Research and Ethics Board (REB 2014-019). Subjects were randomly assigned to one of 3 main groups (eight subjects each). Group 1 subjects tested RAW and 4 subjects from this group were randomly selected to retest using AUG, referred to as subgroup 2. Group 2 subjects tested FLT and 4 subjects from that group were randomly selected to retest using AUG (subgroup 4). Finally, group 3 subjects tested AUG and 4 subjects were selected at random to redo the experiment using RAW (subgroup 5), while the remaining 4 subjects were asked to redo the experiment using FLT, and were referred to as subgroup 6.

Data from Groups 1, 2, and 3 were used to assess differences in outcome measures between the control strategies tested. Additionally, data from subgroups 2, 4, 5, and 6 were used to investigate possible learning effects between the commonly used control strategies and the audio-augmented feedback control strategy (Figure 6). Learning effects between the raw and filtered controllers, subgroups 1 and 3, were investigated in a previous work^35^, which suggested significant improvements in internal model strength and adaptation rate when using the filtered controller after being exposed to the raw controller, but no significant improvements were observed when using the raw controller after being exposed to the filtered controller.

**Figure 6.**
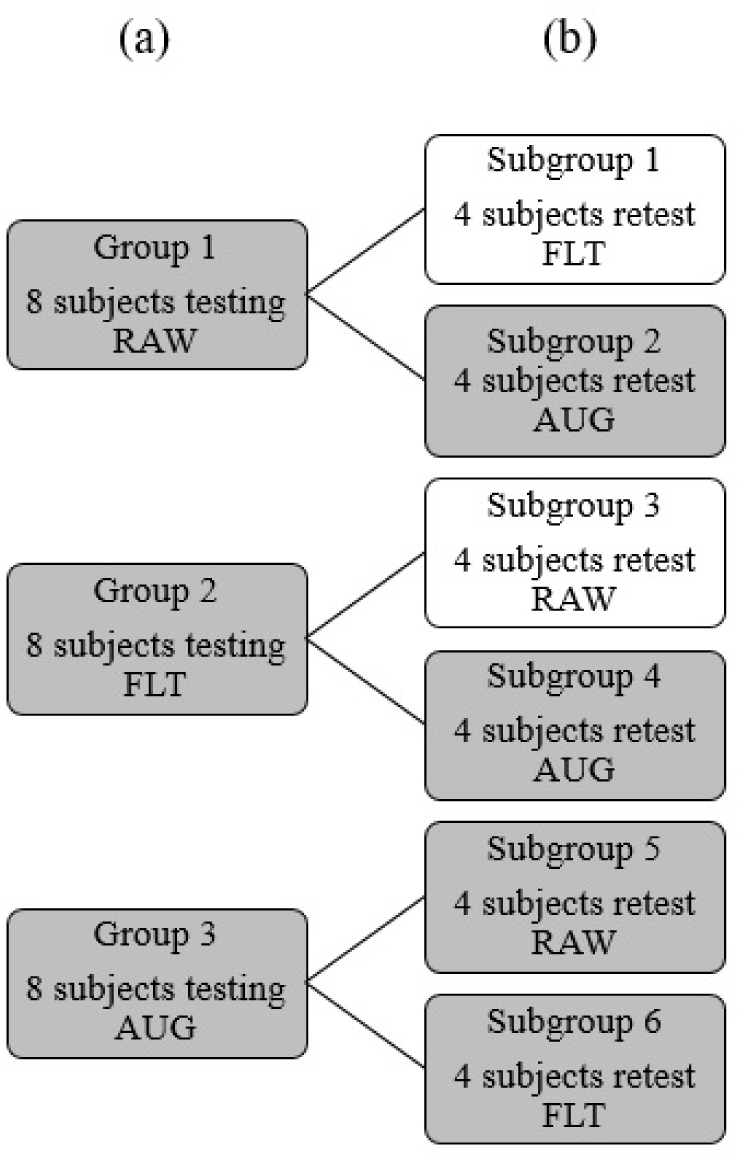
Subject assignment to testing groups. (a) 3 groups of 8 subjects each were used to assess differences in outcome measures between the three control strategies tested. (b) Subgroups of 4 subjects each (in gray) were used to investigate possible learning effects on AUG after using RAW and FLT, and on both RAW and FLT after using AUG.

### Statistical Analysis

The Statistical Package for the Social Science software SPSS (IBM Corp, Released 2016, IBM SPSS Statistics for Windows, Version 24.0. Armonk, NY: IBM Corp) was used to run Levene’s test on JND, adaptation rate, internal model uncertainty, and performance measure results to investigate homogeneity in variances of the data. If data variances were found to be homogenous, one-way ANOVAs were used to assess differences between outcome measures for the control strategies tested. If statistical significance was found, post-hoc analysis was performed using either the Bonferroni or Tukey HSD test and reported using the one with the highest confidence in the p-value^56^. Conversely, if data variances were found to be heterogeneous, robust Welch ANOVA was used instead, and followed by post-hoc analysis using Games-Howell test (Figure 7). Paired two sample t-tests were used with data collected from subjects in the subgroups (equal sample size). All analyses used a significance criterion of *α*=0.05 and error bars shown in data plots were based on the standard error of the mean (SEM)^57^.

**Figure 7.**
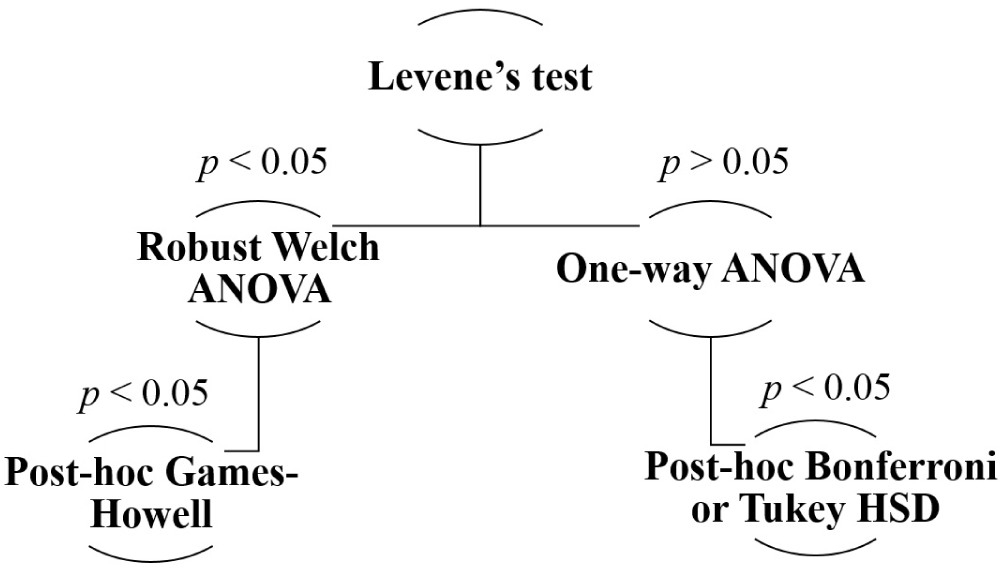
Statistical analysis decision tree followed throughout the analysis of the results in this study.

## Acknowledgements

The authors thank all subjects that volunteered to participate in this study. Many thanks to Dan Blustein for his support and valuable discussions.

## Author contributions statement

A.S., E.S., and J.S. planned the experiments and A.S. prepared and conducted the experiments. All authors analyzed the results, reviewed the manuscript, and approved the submitted version.

## Additional information

**Competing financial interests**: The authors declare that they have no competing interests.

**Data availability**: Data is available upon request – email corresponding author.

